## Supplementary files for "Comprehensive analysis of nasal IgA antibodies induced by intranasal administration of the SARS-CoV-2 spike protein"

**Supplementary Data**

Supplemental Fig.1

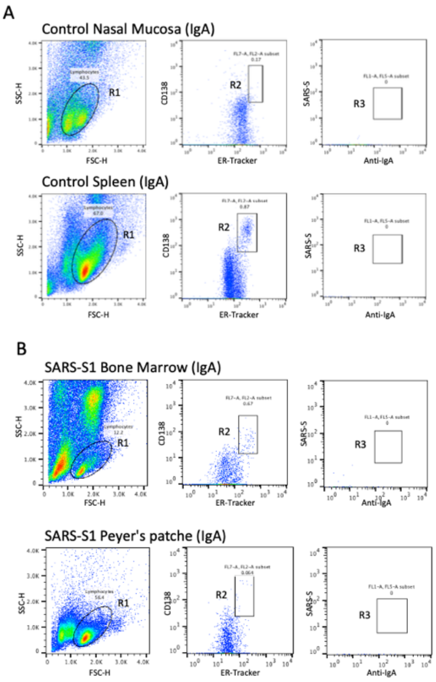


**Supplementary file 1. Intranasal immunization did not induce antigen-specific plasma cells in bone marrow or Peyer's patches.**

(A) Cells isolated from the indicated tissues of PBS-treated mice were treated as shown in Fig. 1B. B

(B) Cells isolated from the indicated tissues of mouse No. 1 were treated as shown in Fig. 1. B. The numbers indicate the percentages of cells in the gated area. A total of 100,000 events were recorded. Representative data are shown.

Supplemental Fig.2

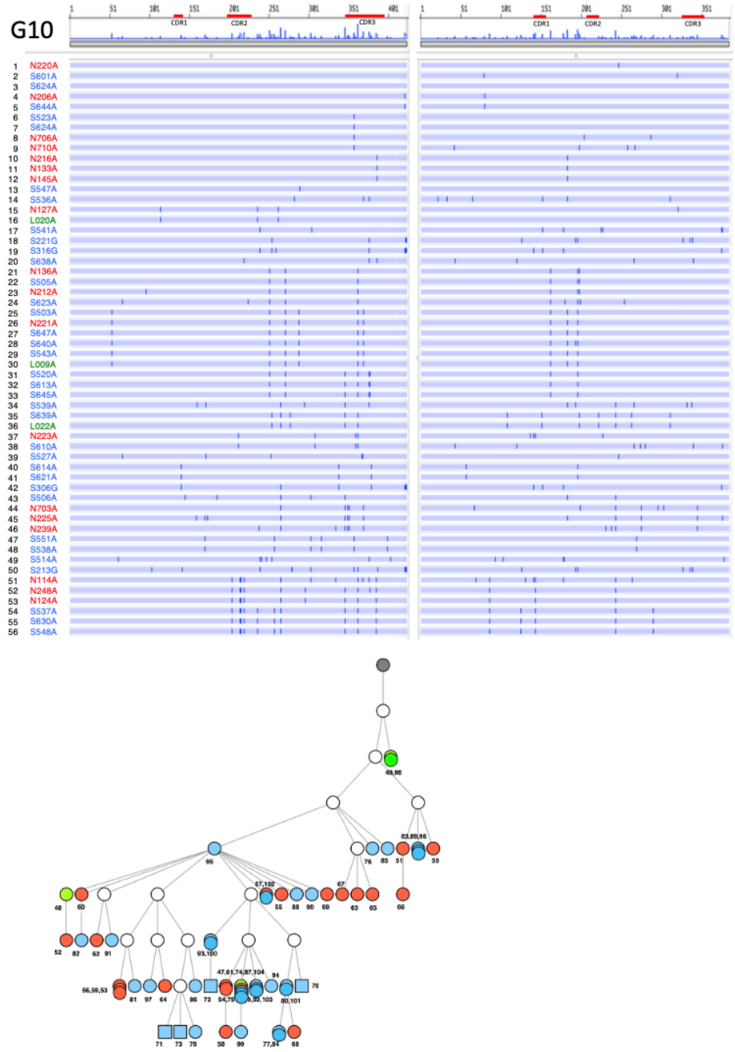


**Supplementary file 2. Nucleotide sequence arraignment of VH and VL genes in the G10 antibodies from No. 1 mouse.**

The VH and VL sequences from the beginning of the signal peptide through the end of FR4 are shown as horizontal lines. Nucleotide changes relative to #N220 are depicted as a vertical bar across the horizontal line. Different colored fonts indicate antibodies derived from the nose (red), spleen (blue) and lung (green). Antibody linage trees based on VH/VK paired sequences are depicted. Gray circles represent the hypothetical germline configuration. White circles represent hypothetical ancestors. Colors indicate nasal (red), splenic (blue) and lung (green) antibodies. Circles and squares indicate IgA and IgG, respectively.

Supplemental Fig.3

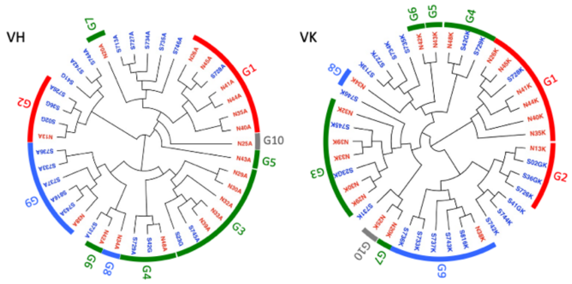


| | | RBD binding | | | | NTD | ACE2-RBD blocking | | | | Neutralize | | | | |
|---|---|---|---|---|---|---|---|---|---|---|---|---|---|---|---|
| Type | Group | Wuhan | Beta | Kappa | Delta | Wuhan | Wuhan | Beta | Kappa | Delta | Wuhan | Nose | Spleen | Total | % of Clones |
| 1 | G1 | | | | | − | | | | | + | 6 | 1 | 7 | 17.1 |
| 1 | G2 | | | | | − | | | | | + | 1 | 3 | 4 | 9.8 |
| 3 | G3 | | | | | − | | | | | − | 5 | 2 | 7 | 17.1 |
| 3 | G4 | | | | | − | | | | | − | 1 | 2 | 3 | 7.3 |
| 3 | G5 | | | | | − | | | | | − | 1 | 0 | 1 | 2.4 |
| 3 | G6 | | | | | − | | | | | − | 1 | 0 | 1 | 2.4 |
| 3 | G7 | | | | | − | | | | | − | 1 | 0 | 1 | 2.4 |
| 4 | G8 | | | | | + | | | | | − | 1 | 0 | 1 | 2.4 |
| 4 | G9 | | | | | + | | | | | − | 1 | 5 | 6 | 14.6 |
| 5 | G10 | | | | | − | | | | | − | 1 | 0 | 1 | 2.4 |
| UN | | ND | ND | ND | ND | ND | ND | ND | ND | ND | ND | 0 | 9 | 9 | 22 |

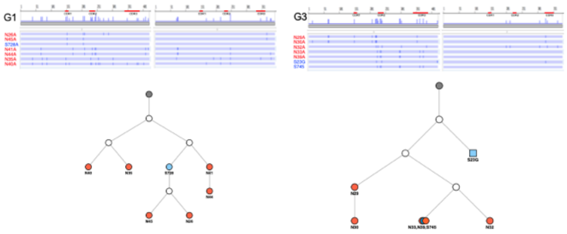


**Supplementary file 3. Characterization of S1-specific monoclonal antibodies obtained from No. 3 mouse**.

(A) A total of 41 S1-reacting antibodies were classified into 17 groups based on their V-(D)-J gene usage. These groups were categorized into five types, as shown in Fig. 1.

(B) Maximum-likelihood phylogenetic tree of the VH and VL chains of the S1-specific monoclonal antibodies. Different colored fonts indicate antibodies obtained from the nose (red) and spleen (blue). Antibody groups are indicated by bands on the outer ring. The color of the band indicates antibody types: Type 1 (red), Type 3 (green), Type 4 (blue) and Type 5 (gray). The prefixes N and S in the antibody clone numbers represent antibodies derived from the nose and spleen, respectively. The suffixes A, G and K in the antibody clone numbers refer to alpha, gamma and kappa chain, respectively.

(C) Nucleotide sequence arraignment of VH and VL genes of G1 and G3 antibodies. The VH and VL sequences from the beginning of the signal peptide through the end of FR4 are shown as horizontal lines. Nucleotide changes relative to N26A and N29A are depicted as vertical bars across the horizontal lines. Different colored fonts indicate antibodies derived from the nose (red) and spleen (blue).

Supplemental Fig.4

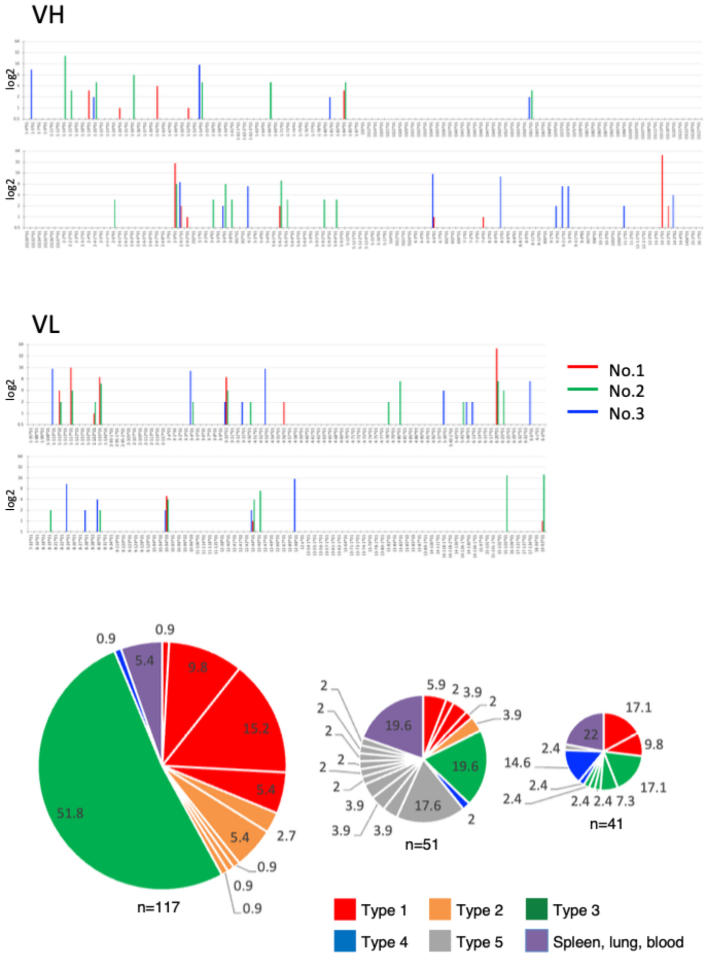


**Supplementary file 4. V-(D)-J gene segment usage in antibodies derived from three immunized mice.**

(A) For each mouse, the numbers of V gene contributions (Y-axis) were determined for each V-gene segment (X-axis). Red, No. 1 mouse; Green, No. 2 mouse; Blue, No. 3 mouse.

(B) The pie chart shows the percentage of antibody types in each mouse. The size of the pie represents the number of antibodies, and slices show the percentage of antibody types.

**Supplementary Table 1 S1-specific antibody sequences obtained from intranasally immunized mice.**

The antibody sequences were compared with the repertoire of mouse germlines compiled at IMGT using IgBLAST (https://www.ncbi.nlm.nih.gov/igblast/). V-(D)-J usage, sequence mismatch and percentage of identities against the highest scoring germline genes as reported in IgBLAST's output are listed. ND, data not determined. The prefixes N, S, L and C in the antibody clone numbers refer to antibodies derived from nose, spleen, lung and blood, respectively. The abbreviations for IgA and IgG are A and G, respectively.

*Please find excel file（Supplementary Table 1）
