## Supplemental Materials for "Comprehensive analysis of nasal IgA antibodies induced by intranasal administration of the SARS-CoV-2 spike protein"

### Supplementary Materials

| Abbreviation | REAGENT or RESOURCE | SOURCE | IDENTIFIER | Working Solution |
|---|---|---|---|---|
| Spike$_{Wuhan}$ | SARS-CoV-2 S protein, His Tag, Super stable trimer | Acro Biosystems, DW, USA | SPN-C52H9 | |
| SpikeWuhan | Biotinylated SARS-CoV-2 S protein (D614G), His,Avitag™, Super stable trimer | Acro Biosystems, DW, USA | SPN-C82E3 | |
| SpikeDelta | Biotinylated SARS-CoV-2 Spike Trimer (T19R, G142D, EF156-157del, R158G, L452R, T478K, D6 | Acro Biosystems, DW, USA | SPN-C82EC | |
| SpikeOmicron | Biotinylated SARS-CoV-2 Spike Trimer, His,Avitag™ (B.1.1.529/Omicron) | Acro Biosystems, DW, USA | SPNC82EE | |
| S1 | Biotinylated SARS-CoV-2 (COVID-19) S1 protein (D614G), His,Avitag | Acro Biosystems, DW, USA | S1N-C82E3 | |
| NTD | Biotinylated SARS-CoV-2 S1 protein NTD, His,Avitag | Acro Biosystems, DW, USA | S1D-C52E2 | |
| RBDWuhan | SARS-CoV-2 (COVID-19) S protein RBD, His Tag | Acro Biosystems, DW, USA | SPD-C52H3 | |
| RBDbeta | SARS-CoV-2 S protein RBD (K417N, E484K, N501Y), His Tag | Acro Biosystems, DW, USA | SPD-C52Hp | |
| RBDkappa | SARS-CoV-2 Spike RBD (L452R, E484Q), His Tag | Acro Biosystems, DW, USA | SPD-C52Hv | |
| RBDdelta | SARS-CoV-2 (COVID-19) S protein RBD (N501Y)), His Tag | Acro Biosystems, DW, USA | SPD-C52HN | |
| ACE2 | Biotinylated Human ACE2 | Sino Biological, Beijing, China | 10108-H08H-B | 5ng/mL |
| CD138 | PE anti-mouse CD138 Antibody | BioLegend, CA, USA | 142503 | 1:250 |
| | Goat Anti-Mouse IgA alpha chain | Abcam, Cambridge, UK | ab97231 | 1μg/50μL |
| | HRP-conjugated goat anti mouse IgA | Abcam, Cambridge, UK | ab97235 | 1:2500 |
| | HRP-conjugated goat anti human IgA | Abcam, Cambridge, UK | ab97215 | 1:2500 |
| | HRP-conjugated goat anti mouse IgG H&L | Abcam, Cambridge, UK | ab6789 | 1:2500 |
| | Streptavidin (HRP) | Abcam, Cambridge, UK | ab7403 | 1:5000 |
| | DyLight-488 labeled goat anti-mouse IgA | Abcam, Cambridge, UK | ab98682 | 1:250 |
| | DyLight-488 labeled goat anti-mouse IgG H&L | Abcam, Cambridge, UK | ab96871 | 1:250 |
| ER-Tracker | ER-Tracke Blue-White DPX | ThermoFisher Scientific, MS, USA | E12353 | 1:1000 |
| | Dynabeads mRNA DIRECT Kit | ThermoFisher Scientific, MS, USA | 61012 | |
| | Nunc MaxiSorp™ flat-bottom | ThermoFisher Scientific, MS, USA | 44-2404-21 | |
| | BluePhos Microwell Substrate Kit | Sera Care | 5120-0059 | |
| | SureBlue/TMB Microwell Peroxidase Substrate | Sera Care | 5120-0076 | |
| | Sensor CHip SA | Cytiva, MS, USA | BR100032 | |
| | Streptavidin Protein, DyLight 650 | ThermoFisher Scientific, MS, USA | 84547 | |
| | CHOgro High Yield Expression System | Takara Baio, Shiga, Japan | MIR6270 | |
| | SARS-CoV-2 direct detection RT–qPCR kit | Takara Baio, Shiga, Japan | RC300A | |
| | Capturem His-Taged Purification Kit | Takara Baio, Shiga, Japan | 635710 | |
| | FreeStyle CHO-S cell | ThermoFisher Scientific, MS, USA | R80007 | |
| | Capturem™ His-Tagged Purification Miniprep Kit | Takara Baio, Tokyo, Japan | 635710 | |
| | Peptide M agarose | ThermoFisher Scientific, MS, USA | gel-pdm-2 | |
| | NativePAGE™ Bis-Tris Gel System | ThermoFisher Scientific, MS, USA | BN1001BOX | |
| | Lympholyte-M | Cedar Lane | CL5030 | |
| | PicaGene Luminescence Kit | Fujifilm Wako | 309-04321 | |
